# Neural excitability increases with axonal resistance between soma and axon initial segment

**DOI:** 10.1101/2020.12.28.424531

**Authors:** Aurélie Fékété, Norbert Ankri, Romain Brette, Dominique Debanne

**Affiliations:** UNIS, UMR 1072 INSERM, Aix-Marseille Université, Marseille, France; Sorbonne Université, INSERM, CNRS, Institut de la Vision, Paris, France

**Keywords:** neuronal excitability, sodium current, axon initial segment

## Abstract

The position of the axon initial segment (AIS) is thought to play a critical role in neuronal excitability. In particular, empirical studies have found correlations between a distal shift in AIS position and a reduction of excitability. Yet, theoretical work has suggested that the neuron should become more excitable as the distance between soma and AIS is increased, because of increased electrical isolation. Specifically, resistive coupling theory predicts that the action potential (AP) threshold decreases with the logarithm of the axial resistance (R_a_) between the middle of the AIS and the soma. However, no direct experimental evidence has been provided so far to support this theoretical prediction. We therefore examined how changes in R_a_ at the axon hillock impact the voltage threshold (V_th_) of the somatic AP in L5 pyramidal neurons. Increasing R_a_ by mechanically pinching the axon between the soma and the AIS was found to lower the spike threshold by ~6 mV. Conversely, decreasing R_a_ by replacing a weakly mobile ion (gluconate) by a highly mobile ion (chloride) elevated the spike threshold. All R_a_-dependent changes in spike threshold could be reproduced in a Hodgkin-Huxley compartmental model. We conclude that in L5 pyramidal neurons, excitability increases with axial resistance, and therefore with a distal shift of the AIS.

## Introduction

In most vertebrate neurons, the action potential (AP) is initiated in the axon initial segment (AIS), a few tens of micrometers away from the soma (Bender and Trussell, 2012; Debanne et al., 2011). Several studies have found that inducing a distal shift of the AIS is accompanied with a reduction of excitability (Grubb and Burrone, 2010; Lezmy et al., 2017; Wefelmeyer et al., 2015). However, these changes in excitability could be due to confounding factors (Kole and Brette, 2018), such as phosphorylation of voltage-gated sodium (Nav) channels (Evans et al., 2015) or regulation of Kv channels (Cudmore et al., 2010; Kirchheim et al., 2013; Kuba et al., 2015). In pyramidal neurons, observational studies found no (Thome et al., 2014) or the opposite (Hamada et al., 2016) correlation between AIS position and excitability. Disentangling the various potential factors is experimentally challenging.

In theory, a distal shift of the AIS could increase the electrical isolation of the AIS, resulting in greater excitability (Baranauskas et al., 2013). Specifically, resistive coupling theory (Brette, 2013; Goethals and Brette, 2020; Telenczuk et al., 2017) predicts that the spike threshold varies as −*k*_*a*_.*log(R*_*a*_), where *k*_*a*_ is the activation slope of Nav channels and *R*_*a*_ is the axial resistance between the soma and the middle of the AIS. However, numerical studies have also occasionally reported an opposite relation, depending on model parameters (Grubb and Burrone, 2010; Gulledge and Bravo, 2016; Lezmy et al., 2017).

We therefore attempted to specifically manipulate the axial resistance R_a_ of L5 pyramidal neurons, thereby changing the electrotonic distance between the soma and AIS and we assessed the impact on the voltage threshold (V_th_) of the somatic AP. We found that increasing R_a_ by mechanically pinching the axon between the soma and the AIS lowered the spike threshold by ~6 mV. Conversely, reducing R_a_ by replacing a weakly mobile ion (gluconate) by a highly mobile ion (chloride) elevated the spike threshold by ~2 mV. All R_a_-dependent changes in spike threshold could be reproduced in a Hodgkin-Huxley compartmental model. These data indicate that the spike threshold of L5 pyramidal neurons is inversely related to the axial resistance R_a_ between the soma and AIS, in agreement with resistive coupling theory. Thus, our results suggest that, contrary to previous suggestions, a distal shift in the AIS position along the axon would result in elevated excitation.

## Results

### Elevating R_a_ by axon pinching lowers spike threshold

L5 pyramidal neurons were recorded in whole-cell configuration of the patch-clamp technique with a pipette filled with Alexa-488 to visualize the axon. Only neurons having an axon emerging from the cell body were used in this study (Hamada et al., 2016; Thome et al., 2014). To test the effects of elevated axial resistance, the pre-AIS region of the axon was pinched with the help of two glass micro-electrodes positioned on each side of the axon. In most of the cases, pinching left the axon intact but a reduction of the transversal section of the axon was observed as a drop of Alexa-488 fluorescence where the pinching occurred (**Suppl. Fig. 1**). Only cells with an axonal length greater than 100 μm and displaying variations in the holding current after pinching that did not exceed ± 70 pA were kept for final analysis.

Axon pinching led to a significant change in spike threshold in half of the cases (7/14 cases, comparison of spike threshold at constant latency over 10 trials before versus after pinching, Mann-Whitney U-test p<0.001; **Fig. 1A-C** & **Suppl. Fig. 2**). In all those significant cases, the spike threshold was hyperpolarised, by about −6 mV (mean: −5.6 ± 1.6 mV, n = 7), up to ~10 mV in 3 cells (**Fig. 1C**). This hyperpolarization of the spike threshold was associated with a reduction in the magnitude of the injected current pulse required to maintain spike latency constant (from 617 ± 74 to 399 ± 74 pA, n = 7, Wilcoxon test, p<0.05; mean reduction: −217 ± 72 pA; **Fig. 1D**), denoting an increase in neuronal excitability. In fact, changes in spike threshold were linearly correlated with changes in current step amplitude (R^2^=0.96, p<0.001; **Fig. 1D**). These effects were related neither to the pinching location nor the axonal length (**Suppl. Fig. 3**). No significant variation in resting membrane potential, holding current, input resistance or latency induced by pinching was observed between the group exhibiting a significant hyperpolarization of the spike threshold and that exhibiting no change in spike threshold (**Suppl. Fig. 3**). Taken together, these results indicate that increasing R_a_ hyperpolarizes the AP threshold measured in the cell body.

**Figure 1.**
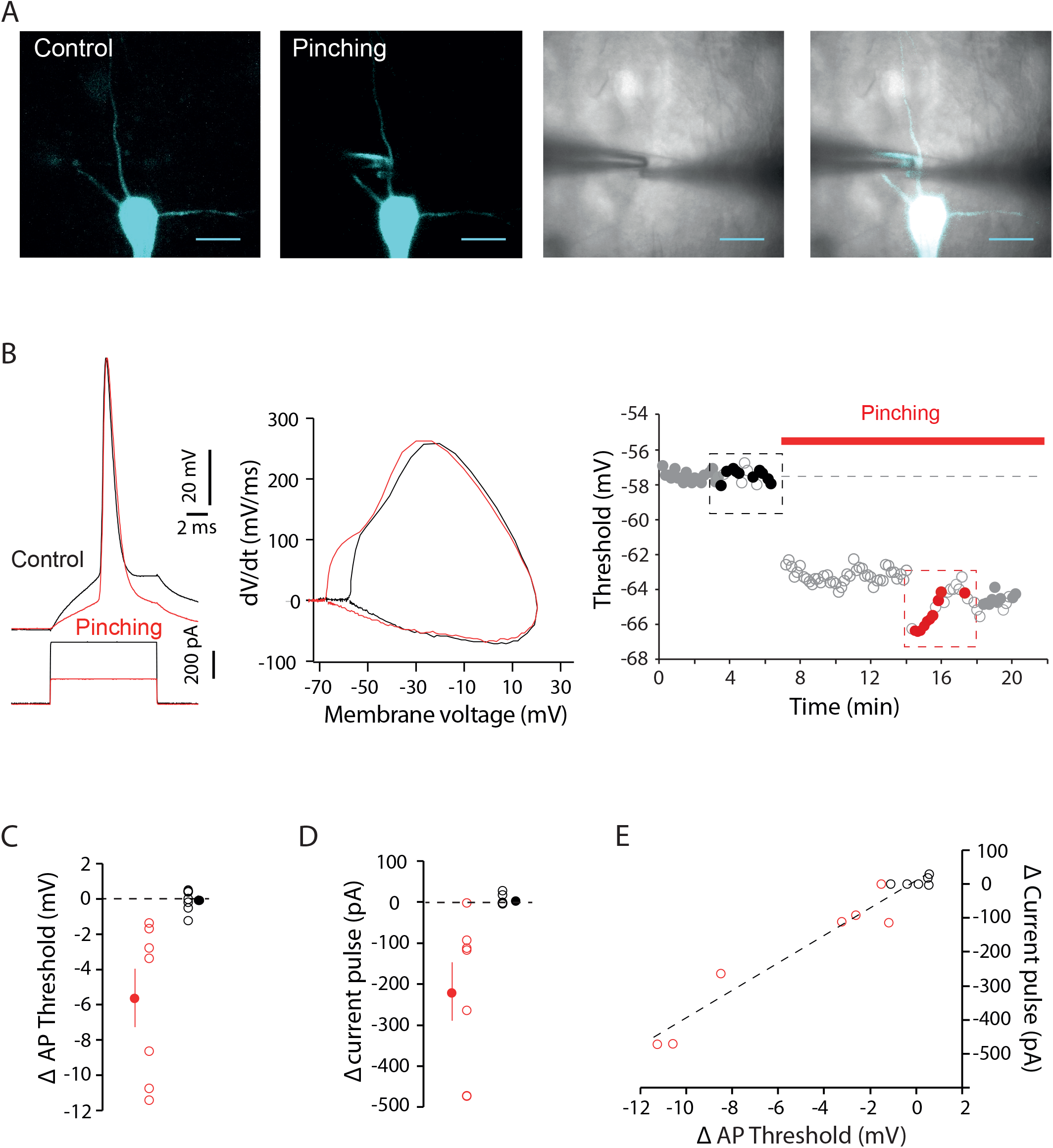
Axon pinching lowers spike threshold. A. Confocal and IR images of the neuron filled with Alexa 488 and imaged before and during axon pinching. Calibration bar: 20 μm. B. Hyperpolarization of the spike threshold during pinching. Left, voltage traces (black, before and red, during pinching). Middle, phase plots. Right, time course of the spike threshold. The effect of pinching on the spike threshold was tested using 10 trials before and during pinching at fixed V_m_ and fixed spike latency (these data are represented in black for the control and in red for the test). The data illustrated as empty gray circles correspond to data that do not meet the selection criteria either because of deviation in membrane potential larger than 1 mV or because of latency variation greater than 1 ms (data in the time period from 7 to 14 min). Here, a significant difference was found between the two samples (Mann-Whitney p<0.001). C. Variation in AP threshold. Neurons showing a significant effect are in red whereas those showing a non-significant effect are in black. Means are represented by filled circles. D. Variation in the current pulse amplitude used to elicit an AP at constant latency. E. Variations in AP threshold were positively correlated with variations in current pulse amplitude (y = (40.62 pA/mV).x + 8.07 pA; R^2^=0.95).

### Dissociation of IS and SD components

Theoretical work suggests that the retro-propagated current corresponding to the initial segment (IS) component varies inversely with R_a_ (Goethals et al., 2020; Hamada et al., 2016). Thus during pinching the current may not be sufficient to trigger a full spike (Michalikova et al., 2017).

In one cell exhibiting a significant hyperpolarization of the spike threshold, the amplitude of the action potential evoked during pinching was found to be considerably reduced in amplitude (**Fig. 2A** and **B**). The presence of this spikelet was not due to any damage caused to the neuron. In fact, a full spike could be recovered when the magnitude of the current step was increased (**Fig. 2C**). This action potential was found to exhibit two different phases clearly visible on the phase plot and identified as the initial segment component and somatodendritic (SD) component (**Fig. 2C**). Interestingly, the slope of the SD component of the AP near the threshold was found to be much slower (6 ms^−1^) compared to the slope of the IS component (21 ms^−1^), suggesting that increasing R_a_ reveals the dissociation of the IS component (fast event) from the SD component (slower event) of the AP.

**Figure 2.**
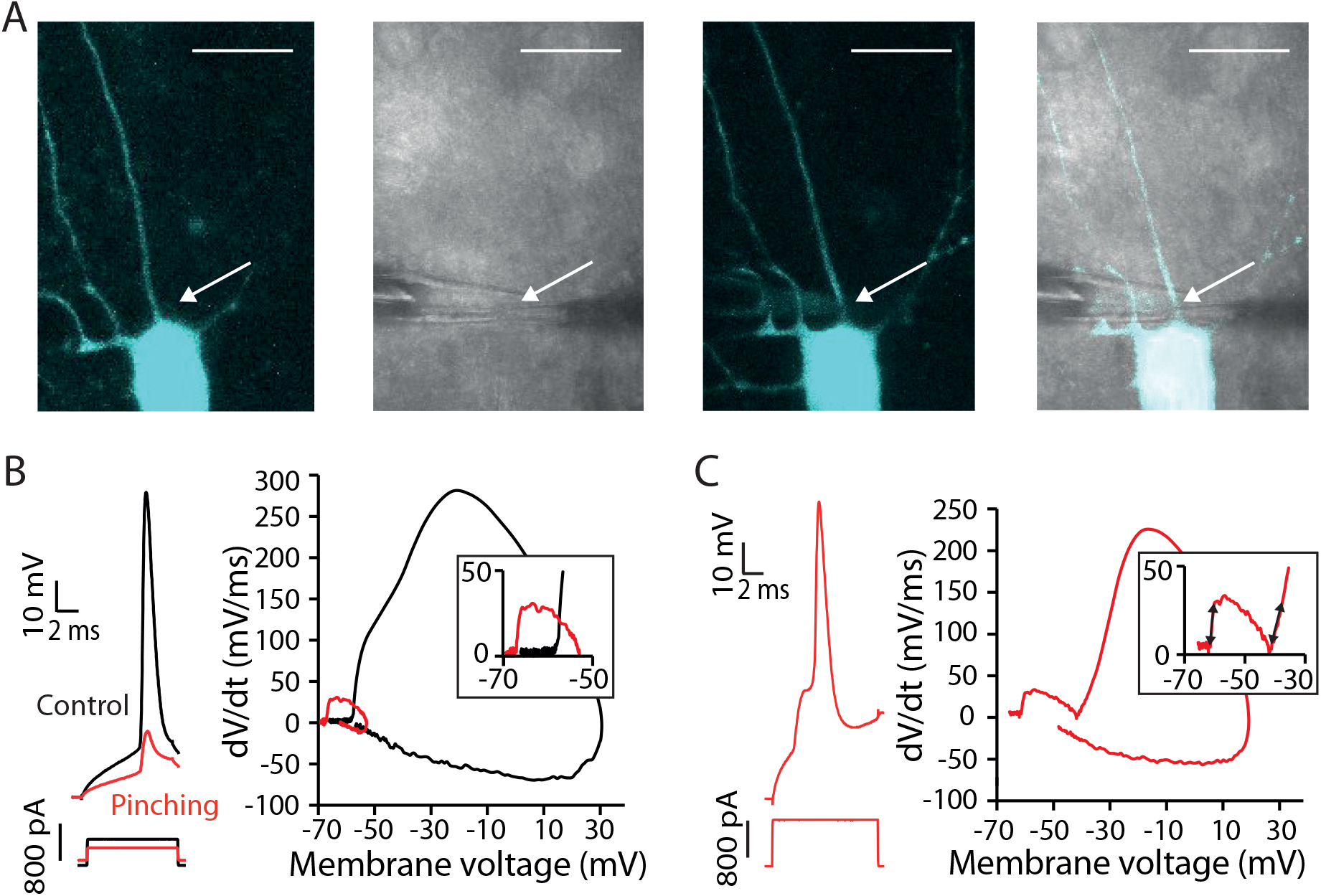
Dissociation of IS from SD component during axon pinching. A. Confocal and IR images of the neuron before and during axon pinching. Calibration bar: 20 μm B. During pinching the spike is hyperpolarized and its amplitude is considerably reduced. The phase plot shows that the spikelet corresponds to the isolated IS component. C. Increasing current amplitude allows recovery of the full spike amplitude corresponding to the SD component. Note the difference in phase slope for the IS component and the SD component in the inset.

In 3 other cases, pinching of the axon accidentally led to ablation of the axon upstream the AIS. This cut was identified by the abrupt interruption of the intra-axonal fluorescence, disappearance of the IS component on the phase plot of the AP, and depolarization of the spike threshold by ~8 mV (from −59.1 ± 0.6 vs. −51.2 ± 2.3 mV; **Fig. 3**). Interestingly, onset rapidness was lower after removal of the AIS (from 37 ± 3 to 14 ± 8 ms^−1^ **Fig. 3C**), indicating that the AP was indeed initiated in the soma rather than in the axon.

**Figure 3.**
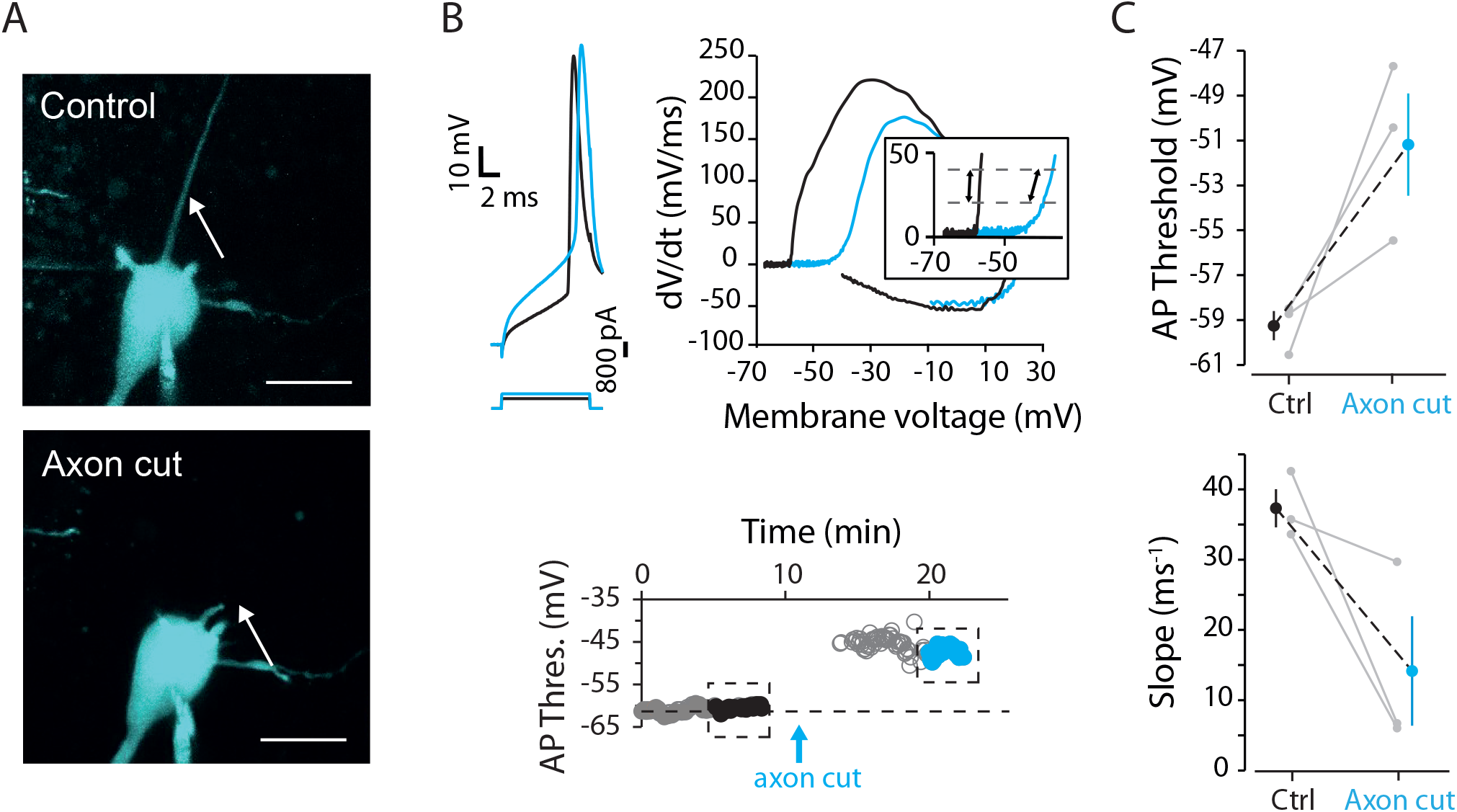
Cutting the axon upstream the AIS elevates AP threshold. A. Confocal images of the neuron before and after axon cut upstream the AIS. Note the cut axon (lower image). Calibration bar: 20 μm. B. Depolarization of the AP threshold. Top left, voltage traces (black before, blue after axon cut). Top right, phase plots. Note the difference in the phase slope in each case. Bottom, time course of AP threshold. The data analyzed in panel C are represented as closed circles (i.e., corresponding to fixed spike delay and constant V_m_). C. Group data of changes in AP threshold and onset rapidness before and after axon cut.

### Lowering R_a_ by intracellular ion substitution elevates spike threshold

To further demonstrate that the spike threshold is inversely related to R_a_, we decreased the intracellular resistivity by replacing a poorly mobile ion (gluconate) by a highly mobile ion (chloride). For this purpose, L5 pyramidal neurons were recorded sequentially with two intracellular solutions of different composition: solution 1, a gluconate-based solution ([KGlu] = 115 mM & [KCl] = 20 mM) and solution 2, a chloride-based solution ([KGlu] = 0 mM & [KCl] = 135 mM). Because gluconate ions are about 3 times less mobile compared to chloride ions (relative mobility of Cl^−^: 1.0388, gluconate ion: 0.33 and K^+^: 1), the total mobility for solution 1 (m_1_) is 194 mM and the total mobility for solution 2 (m_2_) is 275 mM.

The predicted change in spike threshold is given by: 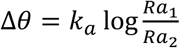. As, R_a_ should be inversely proportional to the total mobility, one predicts that 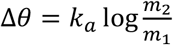. With k_a_ = 6 mV, Δ*θ* = 2.1 mV (the lower threshold is for the solution with the lower mobility, i.e., solution 1).

We next verified experimentally this theoretical prediction (**Fig. 4A**). When L5 pyramidal cells were patched with a gluconate-based solution and then re-patched with a KCl-based solution, the spike threshold was found to be depolarized in all tested cells (n = 6), by ~2 mV on average (1.9 ± 0.6 mV; Wilcoxon test, p<0.05; **Fig. 4B**). No difference in the magnitude of the current pulse was noted after re-patching with KCl-based solution (from 524 ± 74 in K-gluconate to 492 ± 71 pA, n = 6). We conclude that lowering R_a_ by intracellular ion substitution elevates spike threshold in L5 pyramidal neurons, in agreement with the prediction of resistive coupling theory.

**Figure 4.**
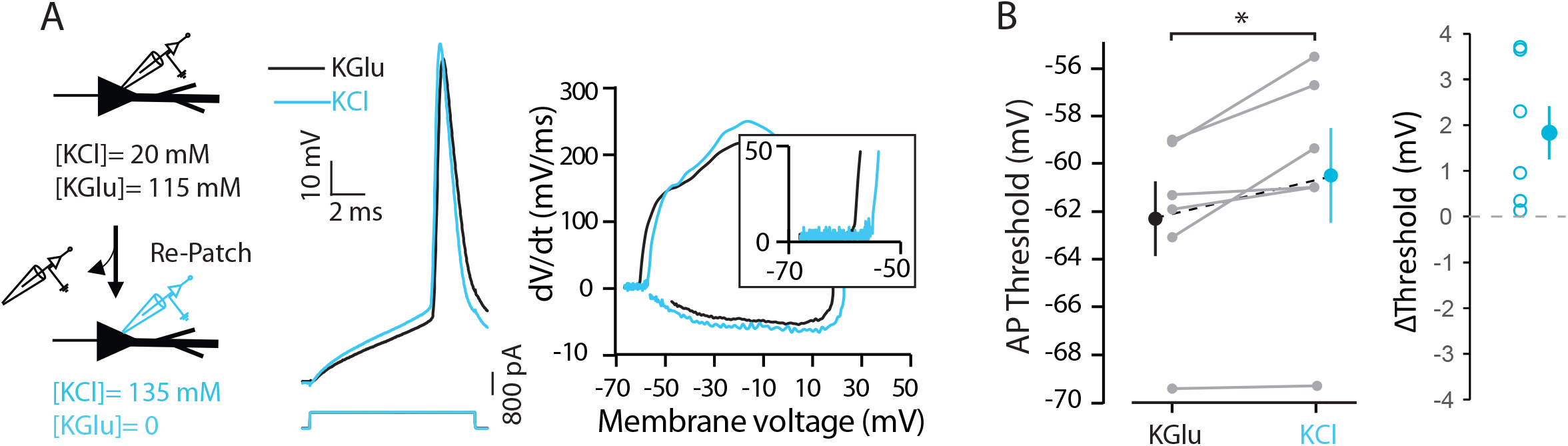
Lowering R_a_ by intracellular ion substitution elevates spike threshold. A. Example of recording a neuron in KGlu and KCl. Note the higher spike threshold on the phase-plot. B. Group data of AP threshold change (*, Wilcoxon, p<0.05).

### Computer modelling of R_a_ change

We reproduced these findings with a compartmental Hodgkin-Huxley model. The model included a cell body, a dendrite and an axon. When the pre-AIS region was pinched (i.e. modelled as a reduction of the diameter from 2.5 μm to 0.48 μm in the pre-AIS region corresponding to an increase in R_a_ from 11 to 50 MΩ between the soma and middle of the AIS, **Suppl. Fig. 4A**), the AP threshold was found to be hyperpolarized by ~5 mV (**Fig. 5A**). The theoretical prediction for this model was k_a_.log(50/11) ~6 mV (Brette, 2013; Goethals and Brette, 2020) (see Methods). Next, we tested the effects of a stronger pinching (corresponding to a further reduction in axon diameter and an elevation in R_a_). As expected, a stronger pinching (i.e. corresponding to a reduction of axon diameter to 0.3 μm and an elevation in R_a_ to 111 MΩ, **Suppl. Fig. 4A**) allowed to isolate the IS component of the spike from the SD component (**Fig. 5B**). As expected, the spike threshold was further hyperpolarized by ~1 mV (**Fig. 5B, 5D**) and the current pulse was reduced (**Fig. 5E**). Next, we simulated the lowering of R_a_ by ion substitution with a uniform reduction in R_i_ by 30% (i.e. the estimated change in resistivity in the experiments). This was found to raise the threshold by ~1.5 mV (**Fig. 5C, 5D**). Thus, the model qualitatively reproduces all the features observed experimentally.

**Figure 5.**
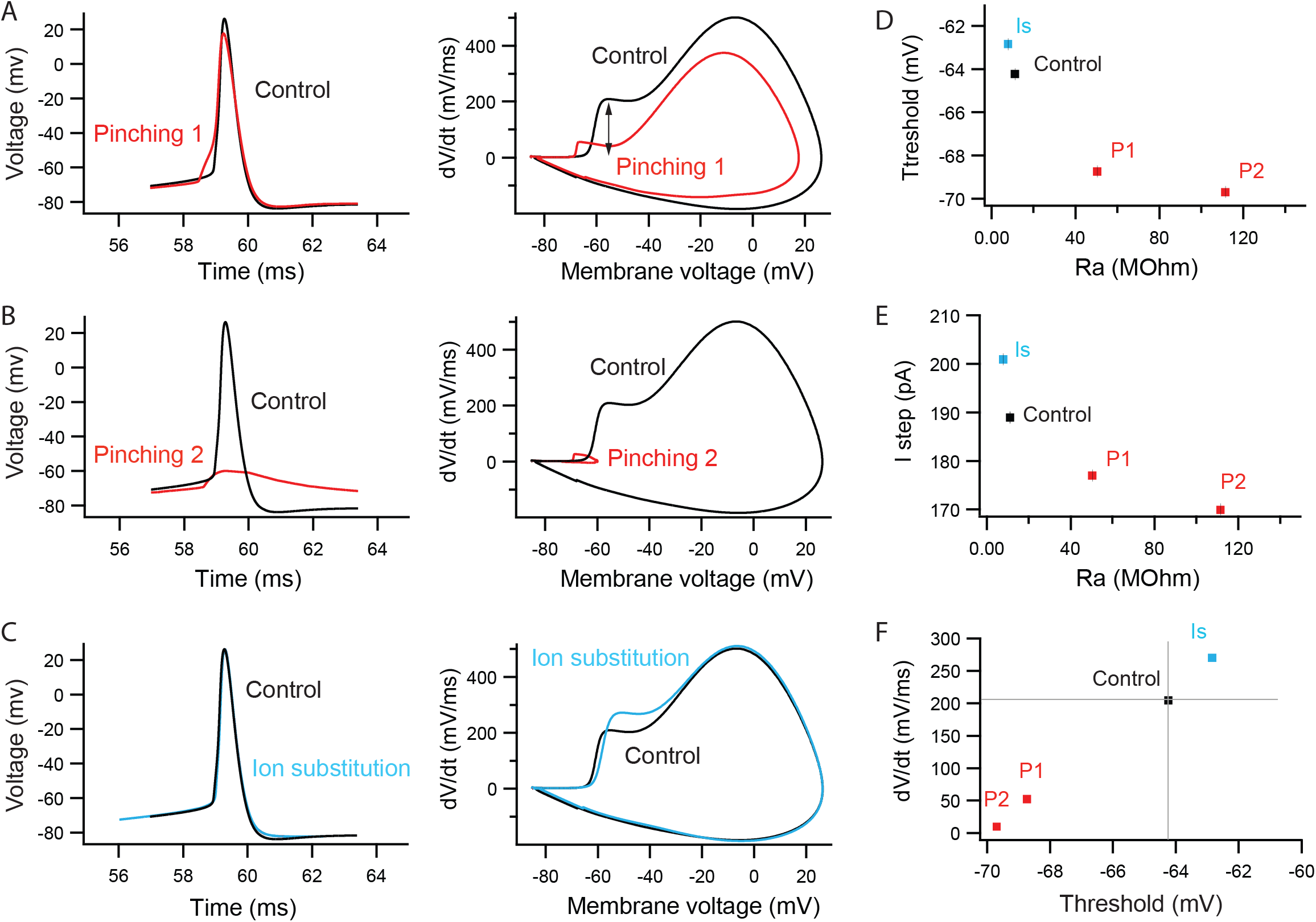
Computer simulation of R_a_ change. A. When R_a_ (taken between the middle of the AIS and the soma) was raised from 11 to 50 MΩ, the spike threshold was found to be hyperpolarized by ~5 mV (see phase-plots). B. Raising R_a_ further to 111 MΩ isolated the IS component (spikelet) and hyperpolarized further the AP threshold (by 1 mV). C. Decreasing intracellular resistivity homogeneously in the neuron to mimic ion substitution raised the spike threshold by ~1.5 mV. D. Plot of AP threshold as a function of R_a_. E. Plot of current pulse amplitude as a function of R_a_. F. Plot of the first maximum of dV/dt in the IS component as a function of threshold.

In accordance with theory (Goethals et al., 2020; Hamada et al., 2016) and previous numerical studies (Telenczuk et al., 2017), in the model the rate of rise of the IS component is reduced when the threshold is hyperpolarized (**Fig. 5F**). We verified that this last relation was also found in the experimental data. The rate of rise of the IS component was measured on the phase plot at V_th_ + 5 mV. A significant correlation was found between the variation in rate of rise of the IS component and the change in AP threshold (R^2^=0.6, p<0.001; **Suppl. Fig. 4B** and **C**). We conclude that modulating the axial resistance at the axon hillock principally affects the IS component measured in the soma.

## Discussion

We have shown here that changes in axial resistance in the pre-AIS region of the axon either by axon hillock pinching or by ion substitution changes the action potential threshold in L5 pyramidal neurons in the opposite direction. First, pinching the axon hillock was found to hyperpolarize the spike threshold by ~6 mV in half of neurons. Because the efficiency of pinching was only visually controlled, the effect was variable from cell to cell. In the remaining cases, no significant effect was observed on the spike threshold (half of cases). In 3 neurons, axon pinching led to an elevation of the spike threshold due to axon cut. Second, replacing a weakly mobile ion (gluconate) by a highly mobile ion (chloride) depolarized the spike threshold by ~2 mV. Finally, a Hodgkin-Huxley compartmental model qualitatively reproduced all the effects observed experimentally. These data suggest that, contrary to previous hypotheses (Grubb and Burrone, 2010; Lezmy et al., 2017), but in agreement with resistive coupling theory (Brette, 2013; Goethals and Brette, 2020; Telenczuk et al., 2017), the specific effect of a distal displacement of the AIS is an increase in excitability.

### Axon hillock pinching

When the axon was intact after axon pinching, we found that AP threshold was hyperpolarized by ~6 mV in a large fraction of the cells. The amplitude of the current that allowed to trigger 1 AP at constant latency was found to be reduced. In fact, the hyperpolarization of the spike threshold was found to be linearly correlated with the reduction of the current pulse, further confirming the increase in excitability when R_a_ was increased by axon pinching.

In one case of pinching, no full spike was recorded in the cell body but instead a spikelet was triggered by the current pulse. The neuron was not damaged since the holding current was unchanged and a full spike could still be evoked upon increase of current step amplitude. Notably, the slope of the phase plot at the spike threshold was found to be much lower for the full spike than for the spikelet, indicating the dissociation of IS and SD components of the AP. These spikelets presumably result from APs that fail to activate sodium channels in the soma (Michalikova et al., 2017). This behaviour was also observed in the neuron model (Fig. 5).

Thus, the scenario of spikelet generation can be reconstituted as follows: axon pinching hyperpolarizes the threshold of the IS spike until a critical value for which the amplitude of the IS spike is too small to activate sodium channels responsible of the SD component and a spikelet is thus induced.

Axon pinching may have mechanically stimulated Kv1 channels that would promote opening of these channels (Hao et al., 2013). This hypothesis is however not plausible for at least two reasons. First, we found no difference in holding current between the group exhibiting a significant threshold hyperpolarization and the other one (**Suppl. Fig. 3**). Second, such mechanosensitive currents would hyperpolarize the AIS, which should theoretically raise the spike threshold at the soma (Goethals and Brette, 2020), the opposite of our observations.

### Axon cut

In a few cases (3 cells), axon pinching led to the ablation of the axon, leaving the cell body isolated from the AIS. Under these conditions, the spike threshold was found to be elevated by ~8 mV and the spike displayed a smooth onset with no kink. Similar depolarizing shift in the spike threshold has been reported upon blockade of Nav channels with local application of TTX on the AIS (Hu et al., 2009; Kole and Stuart, 2008; Palmer and Stuart, 2006) or following genetic disruption of the AIS formation during development (Jenkins et al., 2015; Zonta et al., 2011). In all cases, only somatic Nav channels are functional. The smooth onset of the spike after axon cut indicates that after axon cut, the spike is generated in the cell body.

### Ion substitution

Substitution of gluconate ions by chloride ions was found to elevate the AP threshold by ~2 mV, a value predicted by theory based on ionic mobility. Thus, taken together with the pinching-dependent shift in AP threshold, these data confirm the theory that excitability is inversely related to AIS proximity (Brette, 2013).

### Revisiting the electrophysiological impact of AIS shift

Several studies found that treatments inducing a distal shift of the AIS also resulted in excitability, in contradiction with our findings (Grubb and Burrone, 2010; Lezmy et al., 2017; Wefelmeyer et al., 2015), but observational studies found no such correlation (Hamada et al., 2016; Thome et al., 2014). In both types of study, confounding factors could have contributed to excitability changes (Kole and Brette, 2018), including phosphorylation of Nav channels (Evans et al., 2015) or homeostatic regulation of Kv1 or Kv7 channels (Cudmore et al., 2010; Kirchheim et al., 2013; Kuba et al., 2015). Modelling studies have also shown mixed results, with either a negative (Grubb and Burrone, 2010) or positive (Baranauskas et al., 2013; Brette, 2013) relation between AIS position and excitability, or either case depending on model parameters (Gulledge and Bravo, 2016; Lezmy et al., 2017). Here we have tried to alter the electrotonic distance of the AIS specifically in L5 pyramidal cells, and in all cases we found that an increase in AIS distance was accompanied by an increase in excitability. It remains to be tested whether this relation depends on cell type.

## Materials and methods

### Acute slices of rat neocortex

Neocortical slices (350-400 μm) were obtained from 14- to 20-day-old Wistar rats of both sexes, according to the European and Institutional guidelines (Council Directive 86/609/EEC and French National Research Council and approved by the local health authority (Préfecture des Bouches-du-Rhône, Marseille)). Rats were deeply anesthetized with chloral hydrate (intraperitoneal, 200 mg/kg) and killed by decapitation. Slices were cut in an ice-cold solution containing (mM): 92 *N*-methyl-D-glutamine (NMDG), 30 NaHCO_3_, 25 D-glucose, 10 MgCl_2_, 2.5 KCl, 0.5 CaCl_2_, 1.2 NaH_2_PO_4_, 20 HEPES, 5 sodium ascorbate, 2 thiourea and 3 sodium pyruvate, and were bubbled with 95% O_2_-5% CO_2_, pH 7.4. Slices recovered (1 h) in a solution containing: 125 NaCl, 26 NaHCO_3_, 3 CaCl_2_, 2.5 KCl, 2 MgCl_2_, 0.8 NaH_2_PO_4_ and 10 D-glucose, and were equilibrated with 95% O_2_-5% CO_2_. Each slice was transferred to a submerged chamber mounted on an upright microscope (Olympus BX51WI or Zeiss Axio-Examiner Z1) and neurons were visualized using differential interference contrast infrared videomicroscopy.

### Electrophysiological recordings

Whole-cell recordings from L5 pyramidal neurons were obtained as previously described (Boudkkazi et al., 2007). The external saline contained (in mM): 125 NaCl, 26 NaHCO_3_, 3 CaCl_2_, 2.5 KCl, 2 MgCl_2_, 0.8 NaH_2_PO_4_ and 10 D-glucose, and was equilibrated with 95% O_2_-5% CO_2_. Patch pipettes (5-10 MΩ) were pulled from borosilicate glass and filled with an intracellular solution containing (in mM): 115 K gluconate, 20 KCl, 10 HEPES, 0.5 EGTA, 2 MgCl2, 2 Na2ATP and 0.3 NaGTP (pH=7.4). In experiments where the same neuron was re-patched with a KCl-containing solution, KCl replaced K gluconate ([K gluconate] = 0 mM and [KCl] = 135 mM). Recordings were performed with a MultiClamp-700B (Molecular Devices) at 30°C in a temperature-controlled recording chamber (Luigs & Neumann, Ratingen, Germany). The membrane potential was corrected for the liquid junction potential (−12.2 mV for gluconate-based solution and −2.7 mV for KCl-based solution). Neurons were held at a potential of ~-77 mV. Voltage signals were low-pass filtered (10 kHz), and sequences (200-500 ms) were acquired at 50 kHz with pClamp 10 (Axon Instruments, Molecular Devices).

### Analyses

Electrophysiological signals were analyzed with ClampFit (Axon Instruments). Pooled data are represented as mean ± se in all figures, and we used the Mann-Whitney U-test or Wilcoxon rank-signed test for statistical comparisons. Spikes were evoked by a short (10 ms) current pulse (300-900 pA). As spike threshold is highly sensitive to sodium channel inactivation (Zbili et al., 2020), the latency of the spike was maintained constant (ΔLat<1 ms) before and after axon pinching or axon cut. For this purpose, the amplitude of the current step was adjusted. trials were considered fluctuations in spike latency was maintained within 1 ms. Only trials for which V_m_ was comparable before and after pinching (ΔV_m_<1 mV) were kept for final analysis. For each cell, statistical analysis was performed on the 10 last trials in control vs. the first 10 trials during pinching.

The AP threshold was determined on the spike phase-plot with a custom-made software programmed in LabView. Recording noise was reduced by smoothing the phase-plot with a polynomial of order 4. Then, the AP threshold was determined in the phase plot as the maximum of the second derivative of dV/dt with respect to voltage (Method II described in (Sekerli et al., 2004)).

### Axon pinching and cutting under confocal imaging

To visualize the morphology of L5 pyramidal neurons, Alexa-488 (50 μM) was added to pipette solution and was let to diffuse for at least 10-15 minutes before imaging the axon with a confocal microscope Zeiss (LSM-710) (Rama et al., 2017). Alexa 488 was excited by a laser source at 488 nm. The axon was identified as a thin, aspiny process emerging from the soma or a proximal dendrite at the basal pole of the neuron. Two electrodes were positioned on each side of the axon. To pinch the axon, the two electrodes were gently moved towards each other (Bekkers and Häusser, 2007). The efficacy of pinching was evaluated by visualizing the decrease in fluorescence at the pinched axon. In a few cases, fluorescence was measured with the use of ImageJ (NIH, USA). In a few cases, pinching the axon led to axon cut. In this case, the cut axon was identified by the abrupt disappearance of Alexa fluorescence.

### Hodgkin-Huxley modeling

Computer simulations were performed with LabView 10. The simplified morphology of a L5 pyramidal neuron included dendrite, soma, pre-AIS, AIS and axon. The diameter, length and number of sub-compartments in each neuronal element are provided in **Supplementary Table 1**. The membrane capacitance (C_m_) was set to 0.7 μF/cm^2^ uniformly throughout all compartments (Major et al., 1994). Intracellular resistivity was set to 150 Ω.cm.

The voltage dependence of activation and inactivation of a Hodgkin-Huxley based conductance model (g_Na_, g_Kdr_, g_Kv1_) are given as follows:

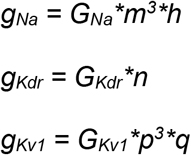

Where *m*, *n* and *p,* are dynamic activation variables for Na, Kdr and Kv1 channels respectively, and *h* and *q* are dynamic inactivation variables for Na current and Kv1 channels, respectively. They evolve according to the following differential equations (Alle and Geiger, 2006):

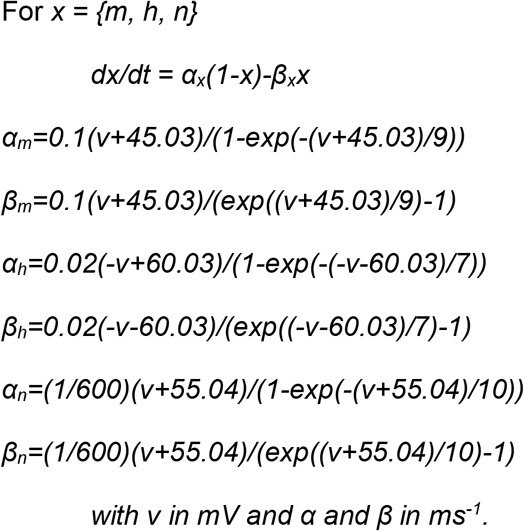

For this model, an asymptotic expansion for hyperpolarized voltages (*ν* → −∞) gives the following exponential scaling for the equilibrium activation variable: *m*_∞_(*ν*)~*e*^*ν*/9 mV^, and thus at spike initiation the sodium current scales with voltage as 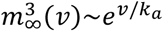 with *k*_*a*_ = 3 mV. This allows us to predict the effect of axon pinching according to the formula *V*_threshold_~ − *k*_*a*_ log *R*_*a*_ (Brette, 2013; Goethals and Brette, 2020).

The Kv1 channel model is taken from (Golomb et al., 2007).

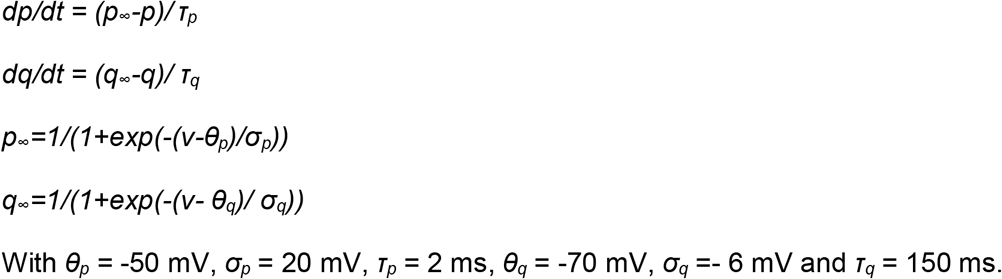

Pinching of the axon by two pipettes was simulated as the narrowing of the axon diameter without change in membrane surface. A piece of axon with a length *L*, resistivity *R*_*i*_ and section *S* has a resistance *R* given by *R* = *R*_*i*_ *L/S*. The axial resistance per unit length *r*_*a*_ = *R*_*i*_/ *S* can be modified equivalently by changing the section S or by changing resistivity from *R*_*i*_ to *R*_*i*_*’* of this element. Thus, changing the diameter from D to D’ can be simulated by changing the resistivity from R_i_ to R_i_’=(D/D’)^2^.

For 3 diameter values of the model, we show in Table 1 the equivalent resistivities and the resulting axial resistances with a reference diameter of 2.5 μm, as illustrated in Suppl. Fig. 4. R_a_ is measured from the soma to the middle of the AIS.

**Table 1:** Intracellular resistivity equivalent to axon diameter changes in the pre-AIS region, and corresponding axial resistances (r_a_: axial resistance per unit length; total R_a_: axial resistance between soma and middle of the AIS). *, current step used to generate a fixed latency AP, see Figure 5E.

| $D_{eq} (\mu\text{m})$ | Resistivity ($\Omega\cdot\text{cm}$) | $r_a$ ($\text{M}\Omega$) | $R_a$ ($\text{M}\Omega$) | $I \text{ step (pA)}^*$ |
| --- | --- | --- | --- | --- |
| 2.5 | 150 | 1.5 | 11.1 | 189 |
| 0.484 | 4000 | 40.7 | 50.3 | 177 |
| 0.306 | 10000 | 101.9 | 111.4 | 170 |

## Acknowledgments

This work was supported by INSERM, CNRS, NeuroMarseille, Agence Nationale de la Recherche (ANR-14-CE13-003 to DD & RB), and the Programme Investissements d’Avenir IHU FOReSIGHT (ANR-18-IAHU-01 to RB).

## Author contributions

AF collected and AF & DD analyzed the data, NA made computer simulations, RB and DD conceived the project, RB, DD and AF designed the experiments, and AF, NA, RB & DD wrote the manuscript.

---

**Supplementary Figure 1.**
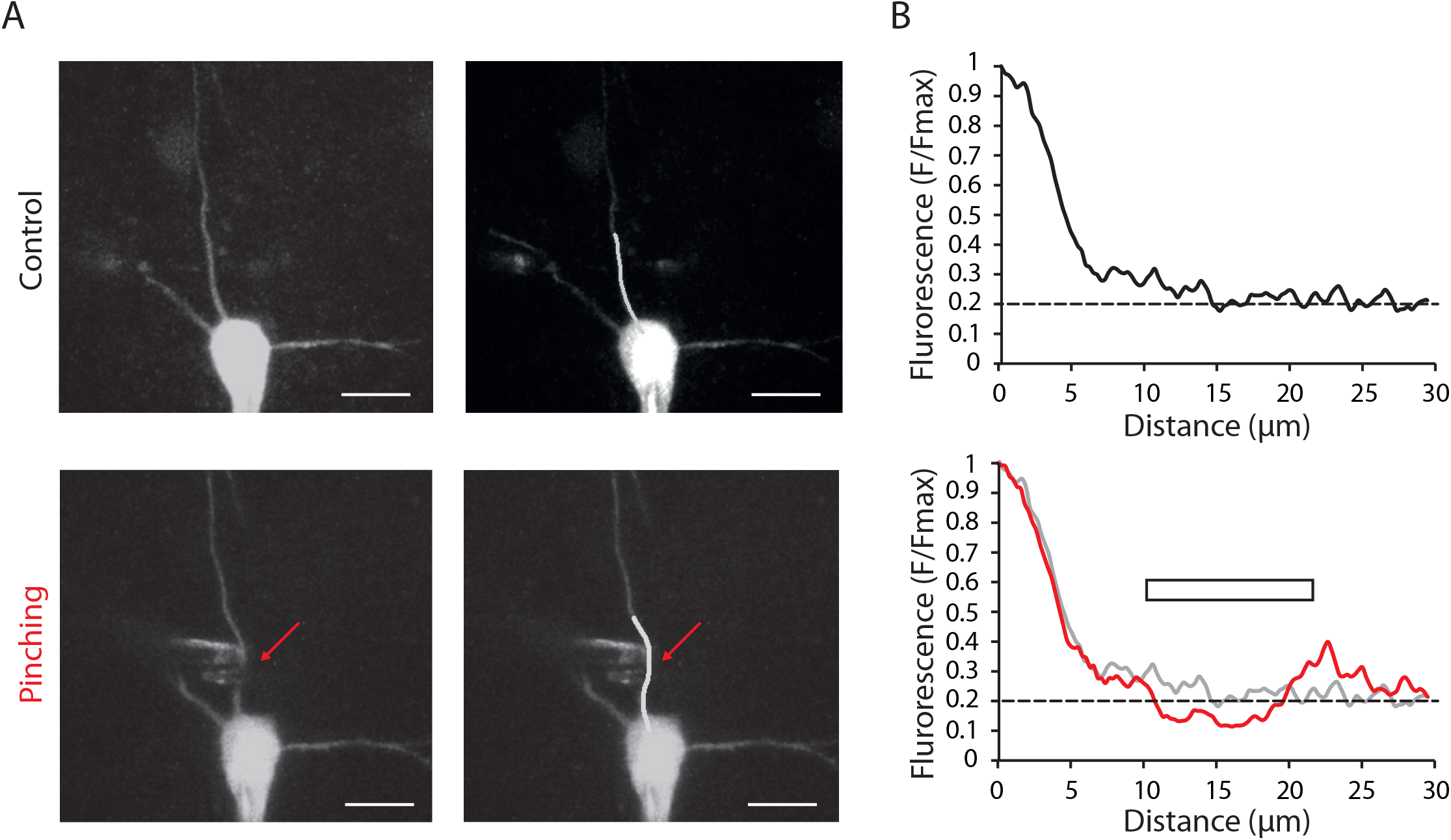
Decrease in intracellular fluorescence caused by the pinching. A. Confocal images of the neuron before and during axon pinching. Calibration bar: 20 μm. Right, image of the neuron with a line materializing the locus of the fluorescence measurement. B. Fluorescence profile of the proximal part of the axon before and during pinching. Note the reduction of fluorescence during pinching (red curve). The horizontal bar indicates the pinching zone.

**Supplementary Figure 2.**
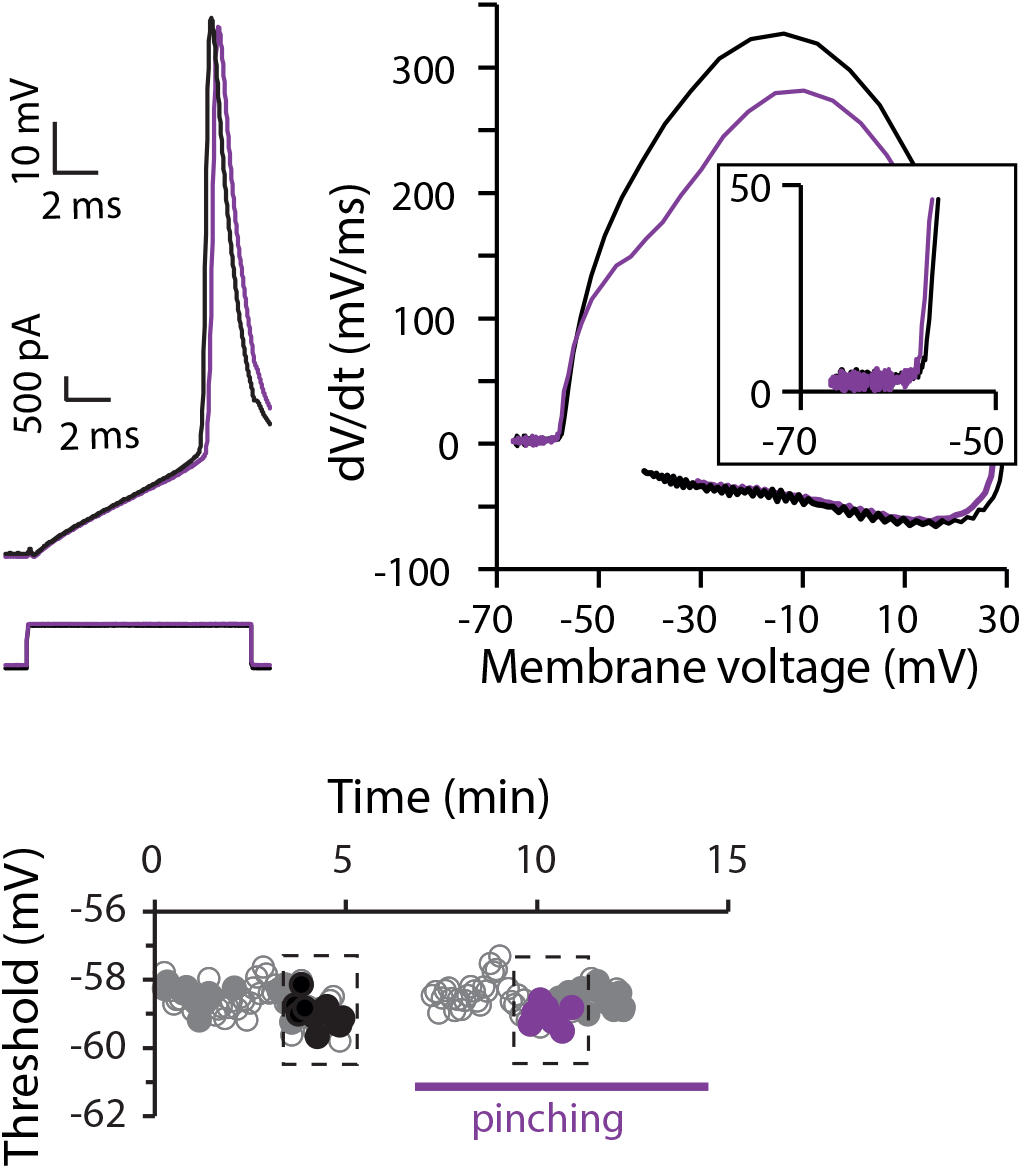
Example of non-significant change in AP threshold induced by axon pinching. A. Voltage traces and phase-plots (as in Fig. 1B). B. Time course of AP threshold. Note the lack of effect on the threshold for the analyzed data illustrated in black and purple (MW, p>0.001).

**Supplementary Figure 3.**
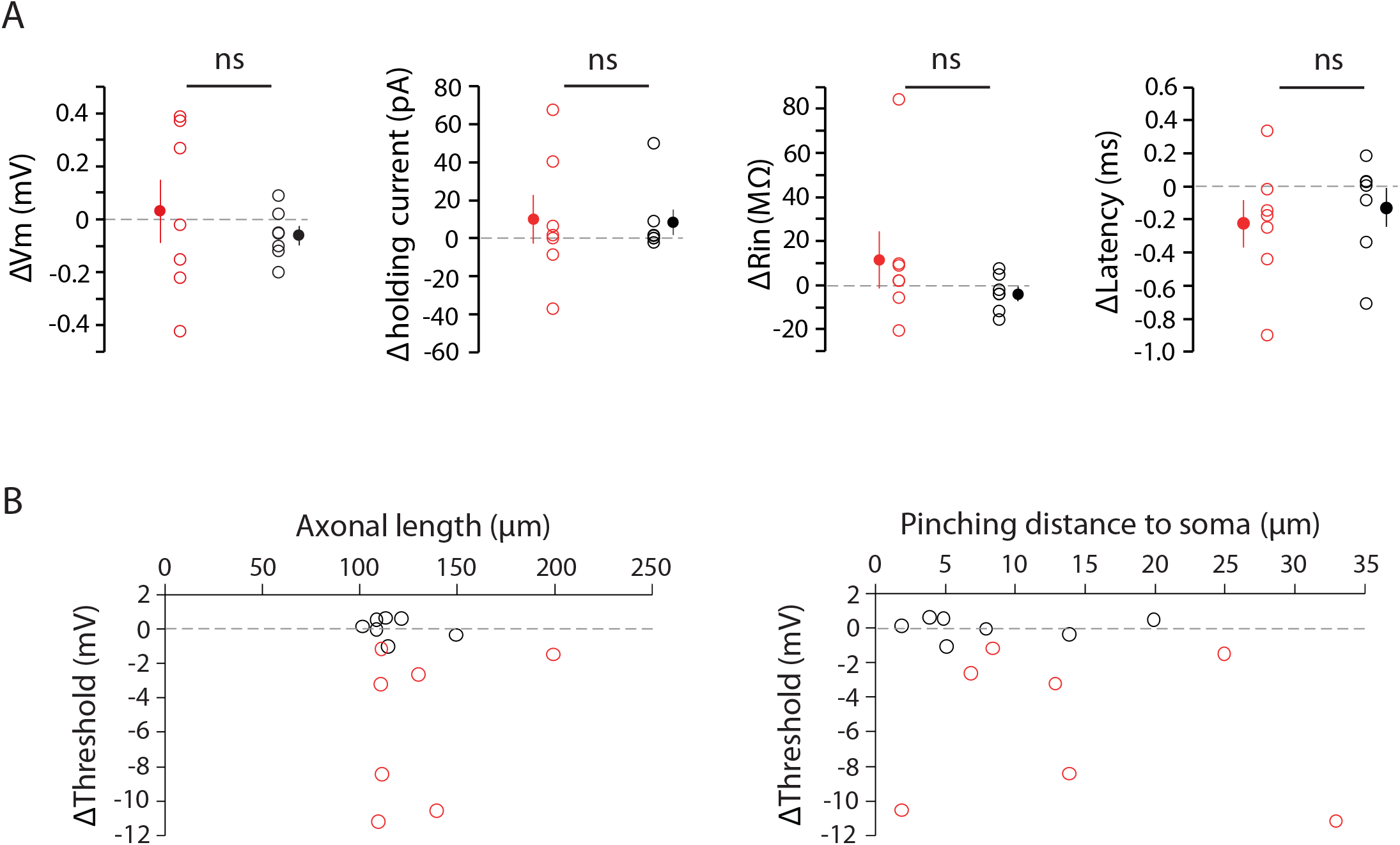
Stability of parameters during pinching. A. Variation in resting membrane potential, holding current, R_in_ and latency during pinching. Neurons showing non-significant effect of pinching on threshold are in black whereas those showing a significant effect are in red. B. Variation in AP threshold as a function of the minimal axonal length and pinching distance.

**Supplementary Figure 4.**
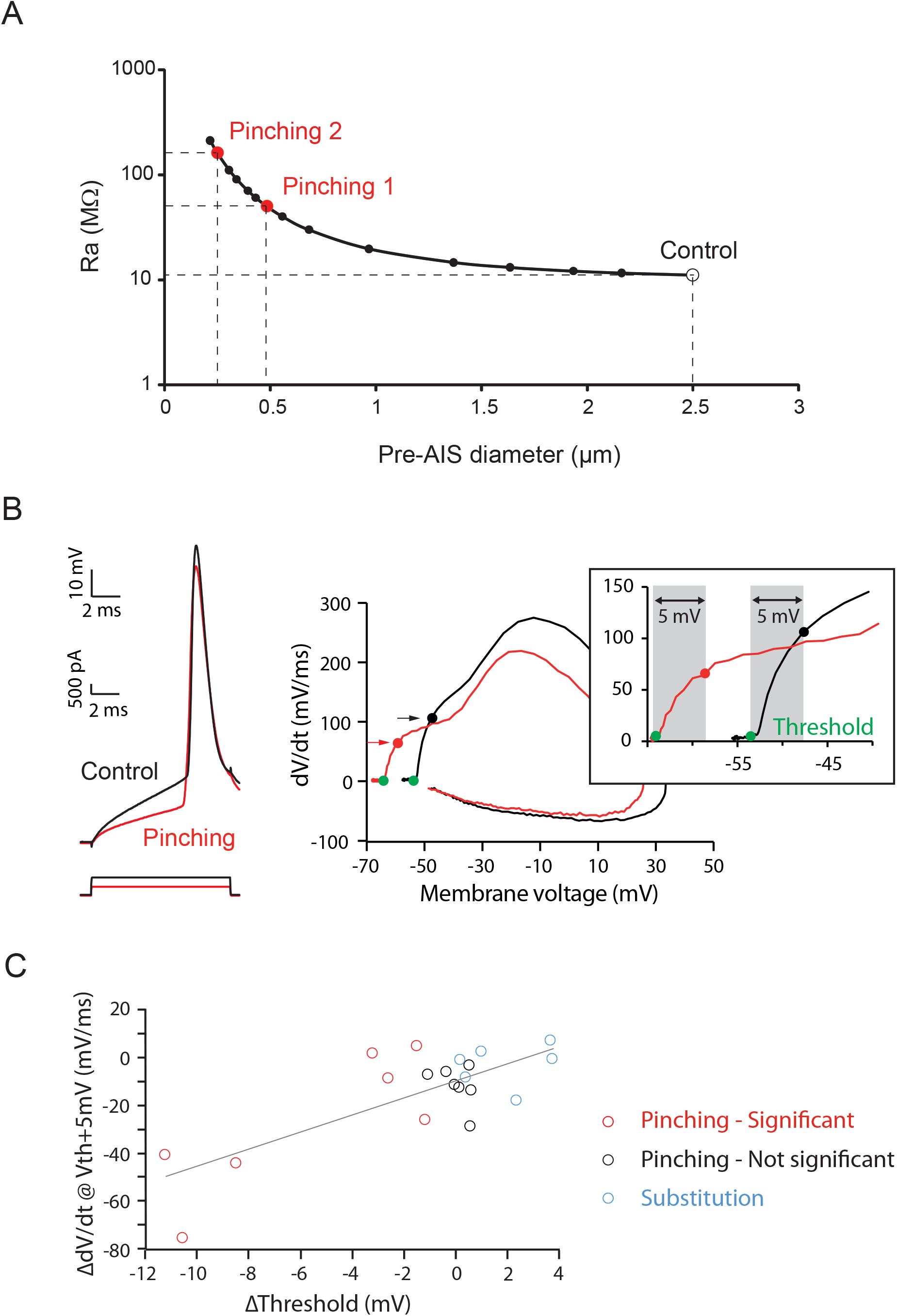
Modeling predictions of axial resistance manipulations. A. Axial resistance (R_a_) between the soma and the middle of the AIS as a function of axon diameter. Note that the ordinate is log scaled. B & C. Experimental verification of the model prediction on the modulation of the rate of rise after pinching and ion substitution. B. Representative example. Left, spikes before and after pinching. Right, phase plot and method for measurements of the rate of rise at threshold + 5 mV. C. Variation in rate of rise as a function of the threshold variation (linear regression y = (3.75/ms)*x - 9.15 mV/ms; p<0.001, R^2^=0.6). Red dots, cells displaying significant change in AP threshold; black dots, non-significant cells and blue dots, cells with substitution of intracellular medium.

**Supplementary Table 1.**
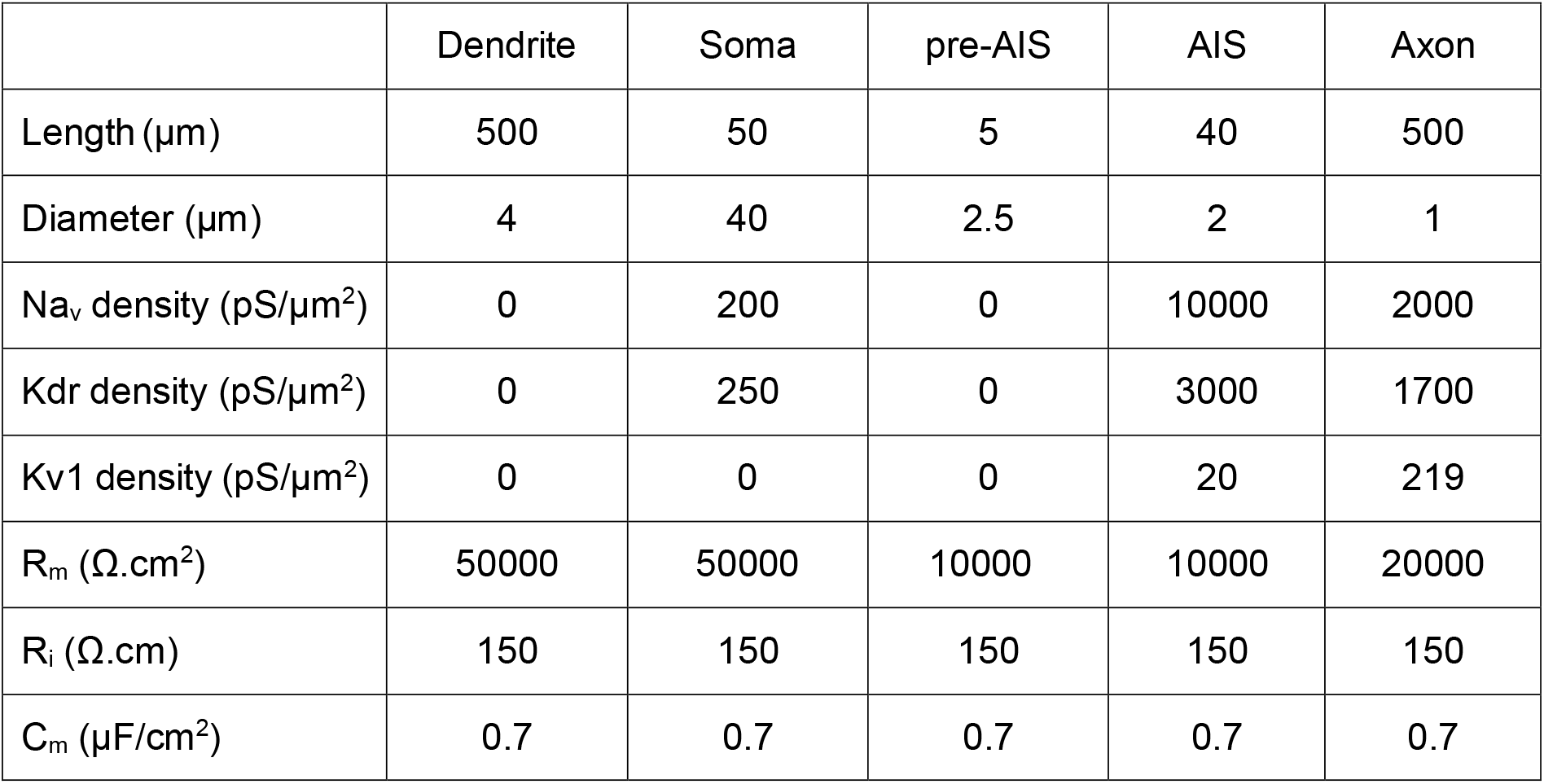
Parameters of the compartmental model

## Notes

### Competing Interest Statement

The authors have declared no competing interest.

